# Selective S1PR1 activation improves brain infarction, neurological outcome, and cerebrovascular endothelial health following experimental ischemic injury

**DOI:** 10.1101/2023.03.27.534410

**Authors:** Samuel X. Shi, Yu-Jing Li, Trevor S. Wendt, Kaibin Shi, Weina Jin, Qiang Liu, Rayna J. Gonzales

**Affiliations:** Department of Basic Medical Sciences, University of Arizona College of Medicine, Phoenix, USA; Department of Neurology, Barrow Neurological Institute, St. Joseph’s Hospital and Medical Center, Phoenix, AZ 85013, USA

**Keywords:** tMCAO, brain endothelial cell, hypoxia plus glucose deprivation, sphingosine-1-phosphate receptor 1.

## Abstract

Sphingosine-1-phosphate receptor 1 (S1PR1) is highly expressed in endothelial cells and receptor activation plays an important role in mediating endothelial function and health, thus showing promise as a pharmacologic target for acute ischemic stroke (AIS) treatment. Here, we examined the effect of a selective S1PR1 ligand, RP101075, on infarct volume and neurological outcome in adult male mice subjected to transient middle cerebral artery occlusion (tMCAO). Concomitantly, we examined S1PR1 expression profile in the ischemic mouse brain, as well as S1PR1 expression and impact of receptor activation on human brain microvascular endothelial cell (HBMEC) proliferation and survival following hypoxia plus glucose deprivation (HGD). We observed that RP101075 administration at onset of reperfusion reduced infarct volume and lessened neurological deficits in tMCAO mice and these responses were S1PR1 dependent. Additionally, we observed that tMCAO increased brain S1PR1 protein levels and flow cytometry revealed increases in S1PR1 levels are greatest in brain endothelial cells compared to other brain cell types (astrocyte, neuron, microglia). In cultured HBMECs, RP101075 increased cell proliferation and ozanimod, parent compound of RP1010175, increased live cell count during HGD; this response was S1PR1 dependent. In conclusion, S1PR1 activation improves neuroprotection/outcome post-stroke and preserves brain endothelial cell survival following ischemia-like injury.

## Introduction

The Food and Drug Administration (FDA) approved treatments for acute ischemic stroke (AIS) include the use of pharmacologic thrombolytics and endovascular mechanical retrieval stents, which provide benefit and improved rates of perfusion.^1, 2^ Unfortunately, their use is time-limited potentiating the risk for cerebral edema formation and increased hemorrhagic transformation. [Reviewed in ^3–5^] Pre-clinical and small-scale clinical pilot stroke studies ^6–10^ have demonstrated the therapeutic efficacy of fingolimod (FTY720), a non-selective sphinogosine-1-phosphate receptor (S1PR) immunomodulator, used in the treatment of relapsing forms of multiple sclerosis (MS). AIS patients receiving fingolimod alone or in conjunction with pharmacologic thrombolysis exhibited a reduction in vascular permeability, reduced neurovascular dysfunction, and decreased neurological deficits within and beyond 4.5 h of ictus.^8, 11^ Although evidence supports fingolimods clinical effectiveness in both MS and stroke patients, this compound has been shown to induce off target effects such as first-dose bradycardia and atrioventricular block ^12, 13^ which could provoke detrimental outcomes in patients with pre-existing heart conditions.

Experimentally, newer generation and more selective S1PR compounds are emerging as promising pharmacologic approaches for AIS therapy. Ozanimod (RPC1063) is a selective S1PR1 and S1PR5 ligand ^14^, and similar to the first generation drug fingolimod, is FDA approved to treat relapsing MS ^15^ and more recently approved to treat ulcerative colitis.^16^ The mechanism of action of long-term administration of ozanimod for these disorders is suggested to work by preventing autoreactive lymphocyte egress to areas of inflammation via S1PR type 1.^14, 17^ Previously, using an in vitro hypoxia plus glucose deprivation (HGD) human brain microvascular endothelial cell (HBMEC) model, we reported that ozanimod decreased levels of the inflammatory mediator, cyclooxygenase-2, following ischemia-like insult.^18^ These data suggest a selective S1PR1 ligand has the potential to elicit a local anti-inflammatory response at the level of the vasculature in the absence of an intact immune response.

In experimental stroke studies, selective S1PR1 ligands demonstrate efficacious outcomes post injury. In two separate experimental mouse stroke models (i.e., intracerebral hemorrhage and transient focal occlusion), ozanimod attenuated blood brain barrier (BBB) permeability, reduced lesion volume, and restored neurological function.^19, 20^ Using an in vivo laser irradiation-induced thrombosis injury model, we reported that RP101075, a novel selective S1PR1 ligand and metabolite of ozanimod, administrated post injury, significantly improved circulation in mouse cerebral arterioles.^21^ Additionally, we reported that RP101075 administration alone had no effect on resting heart. This finding is in line with others that studied ozanimod or similar pharmacologic selective S1PR1 ligands reporting negligible undesirable off target cardiac effects.^16, 17, 22, 23^ Together, these studies suggest that selective S1PR1 ligands elicit protective cerebrovascular effects while potentially negating adverse cardiac effects previously observed with the non-selective S1PR ligand, fingolimod. Since the effect of RP101075 has not been addressed in ischemic stroke, we applied a murine (tMCAO) model and examined whether RP101075 alters infarct volume and neurological outcome post-occlusion and at the onset of reperfusion. We hypothesized that RP101075 elicits neuroprotection and improves neurologic outcome post ischemic injury and we further posited that these responses are S1PR1 dependent.

There are five known S1PR G-protein coupled receptors and are differentially expressed in many cell types. [Reviewed in ^24^] Activation of S1PRs can elicit a variety of biological activities including immune egress via receptor desensitization and potential receptor mediated cell migration, proliferation, and alteration of endothelial function. S1PR1 is coupled with G-protein Gαi ^25, 26^ and receptor activation has been shown to elicit anti-inflammatory properties and promote vascular integrity preservation following stroke or stroke like injury.^18, 27, 28^ In 2016, Scott et al. first identified RP101075 as a metabolite of ozanimod and reported that both ligands exhibit similar S1PR1 selectivity profiles.^29^ Therefore, in the second half of this study, we utilized our in vitro primary HBMEC model to investigate the impact of RP101075 and/or ozanimod (parent compound) on endothelial cell survival following HGD and assessed the involvement of S1PR1 activation. In the brain, S1PR1expression in peripheral immune cells ^30^, neurons ^31^, astrocytes ^32^, microglia ^33^, and endothelial cells ^34–36^ has been well documented. However, the precise S1PR1 expression profile following AIS has not been thoroughly examined. Thusly, we characterized the spatial S1PR1 expression profile in the mouse brain and concomitantly determined changes in S1PR1 levels in cultured HBMECs following ischemic or ischemia-like injury, respectively. By addressing S1PR1 expression and associated selective ligands on mediating cerebrovascular endothelial health and understanding the potential translational impact of these compounds on neurologic outcome post stroke, these studies have further expanded the knowledge base of understanding the role of selective S1PR1 targeting as a therapy for AIS.

## Materials and methods

### Animals

Male C57BL/6 littermate mice (23–25g; 12-wks old; Taconic Biosciences) were housed (12 h light/dark cycle) with free access to food and water. Animal handling and surgical procedures were performed in accordance with recommendations by the Guide for the Care and Use of Laboratory Animals from the National Institutes of Health, and in accordance with the Animal Research: Reporting In Vivo Experiments guidelines. These studies were approved by the Institutional Animal Care and Use Committees at the University of Arizona and Barrow Neurological Institute.

### Transient middle cerebral artery occlusion (tMCAO)

tMCAO was performed using an intraluminal monofilament as previously described.^37–39^ In brief, mice were anesthetized with ketamine (87mg/kg)/xylazine (13mg/kg) mixture (i.p.), ophthalmic lubricant applied to eyes, and body temperature maintained near 37°C during the surgical procedure (Physitemp Instruments LLC., Huron, NJ). An 6-0 nylon monofilament with rounded tip (Beijing Sunbio Biotech Co. Ltd, Beijing China) via right external carotid entry and advanced to internal carotid up through the circle of Willis was used to induce a unilateral focal tMCAO. With the monofilament remaining in place during the occlusion, the neck incision was temporarily closed with suture, and mice were placed in a clean cage on a heating pad to maintain an ambient temperature of 22-23°C at 40% humidity. Following 60 minutes, mice were briefly anesthetized with 3% isoflurane and the monofilament withdrawn at the point of entry. In some experimental groups, W146 ([3R-amino-4-[(3-hexylphenyl)amino]-4-oxobutyl]-phosphonic acid mono trifluoroacetate, 10 μM), a competitive S1PR1 antagonist ^40–42^ or vehicle (5% dimethyl sulfoxide [DMSO]/ 5% Tween in 1x phosphate buffered solution [PBS]) was administered i.p. prior to suture withdrawal. Next, RP101075 (0.6 mg/kg) or vehicle (5% DMSO + 5% Tween 20 in saline) was administered as a single dose i.p. The dosage regimen and administration time window for the optimal analysis of W146 and RP101075 were previously determined.^43^ Cerebral blood flow (CBF) was monitored for an additional 5-10 min before the incision site was permanently closed with 3M Vetbond^TM^. Postoperative incision site pain and discomfort was relieved with lidocaine/prilocaine applied topically. Mice received an injection of warmed saline (0.25mL) subcutaneously for volume replenishment. Anesthesia was discontinued and the mouse was placed back in a clean home cage with moistened food and water for recovery. Sham controls were subjected to the same surgical procedure and post-surgical recovery regimen except for the monofilament placed in the isolated artery. Mice exhibiting residual CBF <20% of pre-ischemic levels throughout the ischemic period and CBF recovery >50% within 10 minutes of reperfusion were included.

### Magnetic resonance imaging

Magnetic resonance imaging (MRI) was used to assess infarct volume 1- and 3-day following tMCAO. Scan acquisitions were performed on a Biospec 7T small animal MRI (Bruker Daltonics Inc., Billerica, MA, USA) as previously described.^38, 44, 45^ Mice were induced with 3-4% isoflurane vaporized using medical grade air maintained at 1.5% via a nose cone, positioned supine on a nonmagnetic warming pad and temperature maintained at between 37.0 +/− 0.5°C. Scans were acquired using Rapid Acquisition Relaxation Enhancement (RARE) sequence with the following parameters: repetition time, TR = 4000ms; effective echo time, TE = 60ms; number of average = 4, field of view (FOV) = 19.2mm×19.2mm, matrix size=192×192, slice thickness = 0.5mm. The MRI data were analyzed with ImageJ software (National Institutes of Health) as previously reported.^38^ Infarct volume was calculated as: infarction volume = lesion volume - (volume of the ipsilateral hemisphere - contralateral hemisphere).

### TTC staining

In a separate cohort of mice, lesion volume was assessed via TTC (2, 3, 5-triphenyltetrazolium chloride) staining in brain slices following 1-day tMCAO. Mice were anesthetized with 4% isoflurane, exsanguinated via a midline cardiac puncture, and whole brain immediately removed, rinsed in 0.9% saline and placed in −20℃ for 15 minutes. Brain slices (2 mm) were consecutively sectioned coronally using a stainless-steel mouse brain matrix (Roboz Surgical Instrument Co., Gaithersburg MD) and stained with 2% TTC for 15 min at 37℃. Next sections were fixed in 4% paraformaldehyde and digital images were obtained and infarct volume quantified using Image-pro plus software (Media Cybernetics, Inc. Rockville, MD): infarction volume = lesion volume - (volume of the ipsilateral hemisphere - volume of the contralateral hemisphere).

### Neurologic assessment

Neurodeficits were evaluated prior to tMCAO and at 1, 3 and 7 days post reperfusion using the modified Neurological Severity Score (mNSS) guidelines, which is comprised of a battery of motor, sensory, reflex and balance tests as previously reported.^37–39^ The rating scale was as follows: a score of 13–18 indicates severe injury, 7–12 indicates moderate injury, and 1–6 indicates mild injury. Each mouse was assessed on a scale from 0 to 18 after recovery from either the tMCAO surgical procedure or sham control. Neurological deficit assessment was performed by an experimenter blinded to experimental groups.

### Flow cytometry

Single-cell suspensions were prepared from ice-cold PBS perfused whole brain (minus the cerebellum) collected 1-day post reperfusion onset and following either tMCAO or sham operation. Tissues were homogenized in ice-cold PBS, digested with collagenase IV (1mg/ml) for 30 minutes at 37°C, then centrifuged (400g for 10 minutes) and the resulting pellet was re-suspended in 30% Percoll (GE Healthcare BioScience AB, Uppsala, Sweden) and centrifuged at 500g for 30 minutes. The resulting cell pellets were washed twice and centrifuged (400g, 8 minutes) in PBS. Pellets were then re-suspended in 1% bovine serum albumin (BSA). Conjugated flow antibodies of EDG-1 (S1PR1), NeuN, CD31(PECAM-1), CD11B, CD45, GFAP were added and incubated at room temperature for 20 minutes (Antibody details provided in Supplemental Materials Table 1). The cells were washed twice with PBS, centrifuged at a low-speed spin (400 g, 8 minutes) and resuspended in PBS for further analysis. Flow cytometric measurements were performed on a FACS Fortessa (BD Biosciences, San Jose, CA) and analyzed using Flowjo 7.6 software (Informer Technologies, Ashland, OR).

### HBMEC culture and hypoxia plus glucose deprivation models

Male primary human brain microvascular endothelial cells (HBMECs) purchased (Cell Systems; Kirkland, WA, Catalogue number: ACBRI 376 Lot number: 376.05.02.01.2F) cryopreserved subcultures at passage 3 were grown in a 5% CO_2_ incubator at 37°C in phenol red-free Complete Classic Medium (10% FBS; fetal bovine serum; Catalogue number: 4Z0-500) with Bac-off Antibiotic (Cell Systems, Catalogue number: 4Z0-643), and CultureBoost™ (Cell systems, Catalogue number: 4CB-500). Cells were seeded at a density of 4 to 5×10^5^/cm^2^ and reached 80% confluence within 5 to 7 days. At that time, designated plates were cryopreserved for future use or continued in culture and studied at passage 6-7. Von Willebrand factor, PECAM-1, and endothelial nitric oxide synthase protein expression validated endothelial origin (data not included). Culture medium for HGD was replaced with glucose-free DMEM, 10% FBS, Bac-off Antibiotic, CultureBoost™. Medium in designated normoxic plates was replaced with fresh glucose containing DMEM spiked with 10% FBS, Bac-off Antibiotic, and CultureBoost™. In cultured exposed to HGD alone, prior to exposure, plates were treated with vehicle, an S1PR1 ligand, S1PR1 antagonist or S1PR1 antagonist plus ligand. The HGD plates were placed in plexiglass incubator sub-chamber (C-Chamber Bio-Spherix; Lacona, NY) housed within a 5% CO_2_ incubator at 37°C. The hypoxic chamber was flushed with medical grade gas mixture of 1% O_2_, 5% CO_2_ and nitrogen balance and exposed for 3h. In cultured exposed to HGD with reperfusion (HGD/R), immediately following 3h HGD exposure, medium containing glucose was returned to the plates followed by immediate treatment with vehicle, an S1PR1 ligand, S1PR1 antagonist or S1PR1 antagonist plus ligand and returned to 5% CO_2_ incubator at 37°C and exposed to room air for 12h or 24h. Normoxia groups remained within a separate 5% CO_2_ incubator at 37°C and exposed to room air.

### Western blot

Tissue/cell lysate preparation and standard SDS-PAGE were performed as previously described.^27^ Methodological details are provided in the Supplemental Materials Methods section and antibody details provided in Supplemental Materials Table 2.

### Crystal violet staining

Endothelial cell vacuolization and vesicles were assessed using crystal violet staining following exposure to either normoxia or HGD. HBMECs were seeded in 6-well plates and at 75% confluency were stained for 20 min at RT using a crystal violet (Fisher Scientific) fixing (37% formaldehyde) /staining buffer. Stained HBMECs were then gently washed with nanopure water for excess crystal violet stain and images captured with inverted digital microscope (Keyence; Catalogue number: BZ-X800). Cell vesicle count was assessed using FIJI (NIH; enhanced version of ImageJ2) images from the crystal violet experiments as previously described ^27^ and modified slightly to selectively quantitate smaller vesicle structures in this study.

### Proliferation analysis

A BrdU (5′-bromo-2′-deoxyuridine; 40uM) staining protocol was used to identify proliferating cells following either normoxia or HGD exposure and drug treatment. HBMECs were seeded (1×10^6^/well) in 6-well plates in a total volume of 3mL of medium and collected after exposure/drug treatment via trypsin and trypsin-neutralizer and according to manufacturer protocol using FITC anti-BrdU antibody (Biolegend, San Diego, CA). Flow cytometric measurements were performed on a FACS Fortessa (BD Biosciences, San Jose, CA) and analyzed using FACS Diva and FlowJo 7.6 software (Informer Technologies, Ashland, OR).

### Live cell count

To determine the number of viable cells following exposure to either normoxia or HGD, HBMECs were cultured in 6-well plates and seeded overnight under normoxic conditions in a 5% CO_2_ incubator at 37°C. Next, culture medium was changed and following either normoxia or HGD exposure ±RP101075 or ±ozanimod ±W146 treatment for 3h, cells were collected using trypsin and trypsin-neutralizer and centrifuged at a low-speed spin (400g, 8 min). The resulting trypsin/trypsin-neutralizer supernatant was aspirated, and the remaining cell pellet was resuspended with 100ul of a 0.4% trypan blue solution (Corning) to stain non-viable cells. Trypan blue cell suspension solution (10ul) was taken out and applied to the Reichert Bright-Line metallized hemocytometer (Hausser Scientific). The chamber underneath the coverslip was gently filled, allowing the cell suspension to be drawn out by capillary action. Using an Olympus CKX41 inverted light microscope and 10X objective, the total cell counts were obtained by a blinded analyzer in all 4 sets of 16 squares from which an average value was recorded. The live cell counts were obtained by subtracting the total cell counts by the number of dead cells in all 4 sets of 16 squares and from which an average value was recorded. To calculate the number of live cells per milliliter, the average of the live cell count was multiplied by 10^4^.

### Drug preparation

Prior to each experiment, stock solutions of RP101075 (Receptos Inc) were freshly prepared on the same day of treatments/experiments under sterile conditions as previously described. ^39^ Details for drug preparation are described in the Supplemental Material Methods section.

### Reagents

Reagents were purchased from Sigma-Aldrich Corporation (St. Louis, MO) unless otherwise noted.

### Statistical analysis

Experiments were repeated for statistical analysis using GraphPad Prism 8.0 software (GraphPad Software Inc., La Jolla, CA). A Shapiro-Wilk test confirmed if data sets achieved a normal distribution. An *F*-test confirmed if the data sets achieved equal variance amongst groups. For sets that achieved a normal distribution, a direct comparison between two groups were made using a two-tailed unpaired Student t-test with Welch’s correction. Comparisons between three or more groups were made using either one-way analysis of variance (ANOVA) followed by Tukey post-hoc test, or two-way ANOVA accompanied by Tukey post-hoc test for multiple comparisons. Western analysis consisted of running lysates from each treatment group, including vehicle, on the same blot for direct comparison. Band densities were normalized to loading control, β-actin, and data expressed as fold change relative to control group (i.e., normoxia, normoxia plus vehicle, sham, or sham plus vehicle). Significance was set at α < 0.05. Data are shown as Means ± SD.

## Results

### RP101075 administration post-occlusion and at the onset of reperfusion reduced infarct volume

To determine the effect of RP101075 on brain injury, MRI was used to quantify lesion volume at day 1 and 3 in cohorts of mice that underwent tMCAO and administration a single dose of RP101075 or vehicle at the onset of reperfusion. Representative brain MRI images are illustrated in Figure 1a. Infarct volumes were quantified with ImageJ software and plotted as infarct volume (mm^3^) (Figure 1b). There was no difference in infarct volume in tMCAO mice receiving vehicle at day 1 and at day 3. However, the group that received RP101075 elicited a significant decrease in infarct volume at day 1 and 3 compared to vehicle with the greatest reduction in lesion volume observed at day 3 post-occlusion. These data suggest that RP101075 administration post occlusion and at onset of reperfusion temporally reduced injury in a male mouse model of tMCAO.

**Figure 1.**
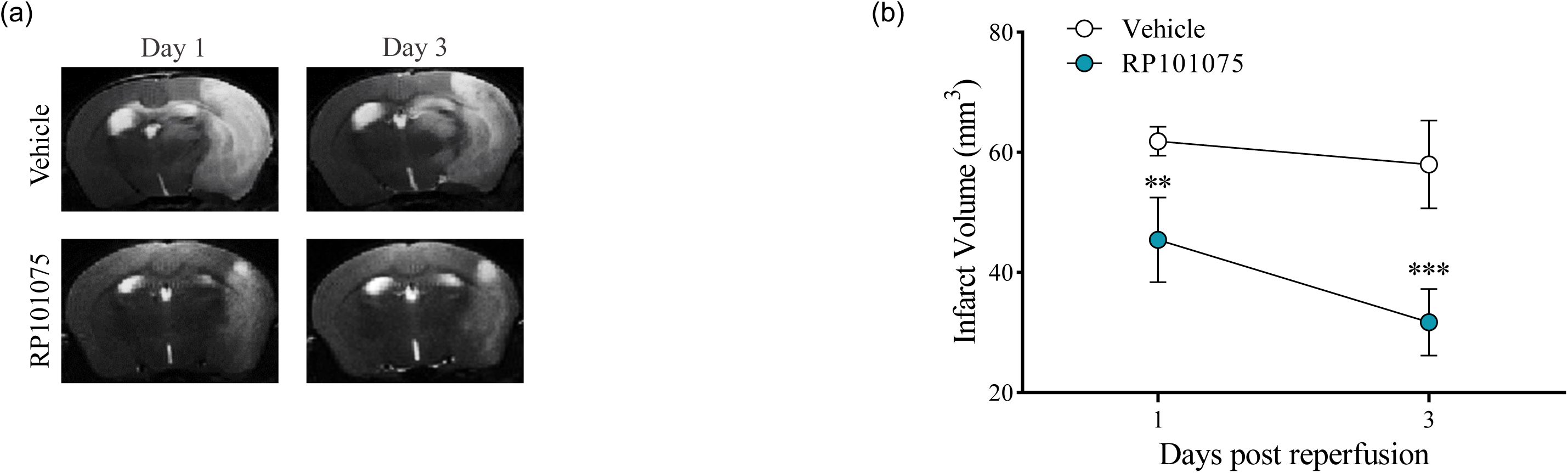
**Selective S1PR1 ligand, RP101075, decreased infarct volume following tMCAO.** (a) Representative 7T Bruker coronal MRI images at day 1 and at day 3 post transient ischemic injury from RP101075 (0.6mg/kg) and vehicle treated groups. (b) Quantification of infarct volume assessed at indicated time points after ischemia and during reperfusion. Data are presented as mean ± SD; N = 5-6/group. Infarct volume by two-way ANOVA with Tukey post-hoc test. Day 1 Vehicle vs Day 1 RP101075 \*\**P* = 0.0014, Day 3 Vehicle vs Day 3 RP101075 \*\*\**P* = 0.0003.

### RP1010175-induced decrease in brain infarct volume following tMCAO is S1PR1 dependent

To determine if the beneficial actions of RP101075 on reducing infarct volume within a transient ischemia/reperfusion mouse model are S1PR1 dependent, tMCAO mice were treated with vehicle or RP101075 in the presence or absence of W146, a selective S1PR1 antagonist. Photographs of representative TTC stained slices 1-day post reperfusion from whole brain isolated from tMCAO mice are shown in Figure 2a. Figure 2b illustrates the data supporting that S1PR1 pharmacologic inhibition attenuated the RP101075 mediated reduction in lesion volume post occlusion and during reperfusion, RP101075+W146 vs. vehicle, 45.803mm^3^ ± 6.147mm^3^ vs. 60.594mm^3^ ± 4.215mm^3^ (P=0.2367). W146 alone had no effect on the infarct volume in tMCAO mice. These data suggest that the reduction elicited by RP1010175 on brain injury post tMCAO is S1PR1 dependent.

**Figure 2.**
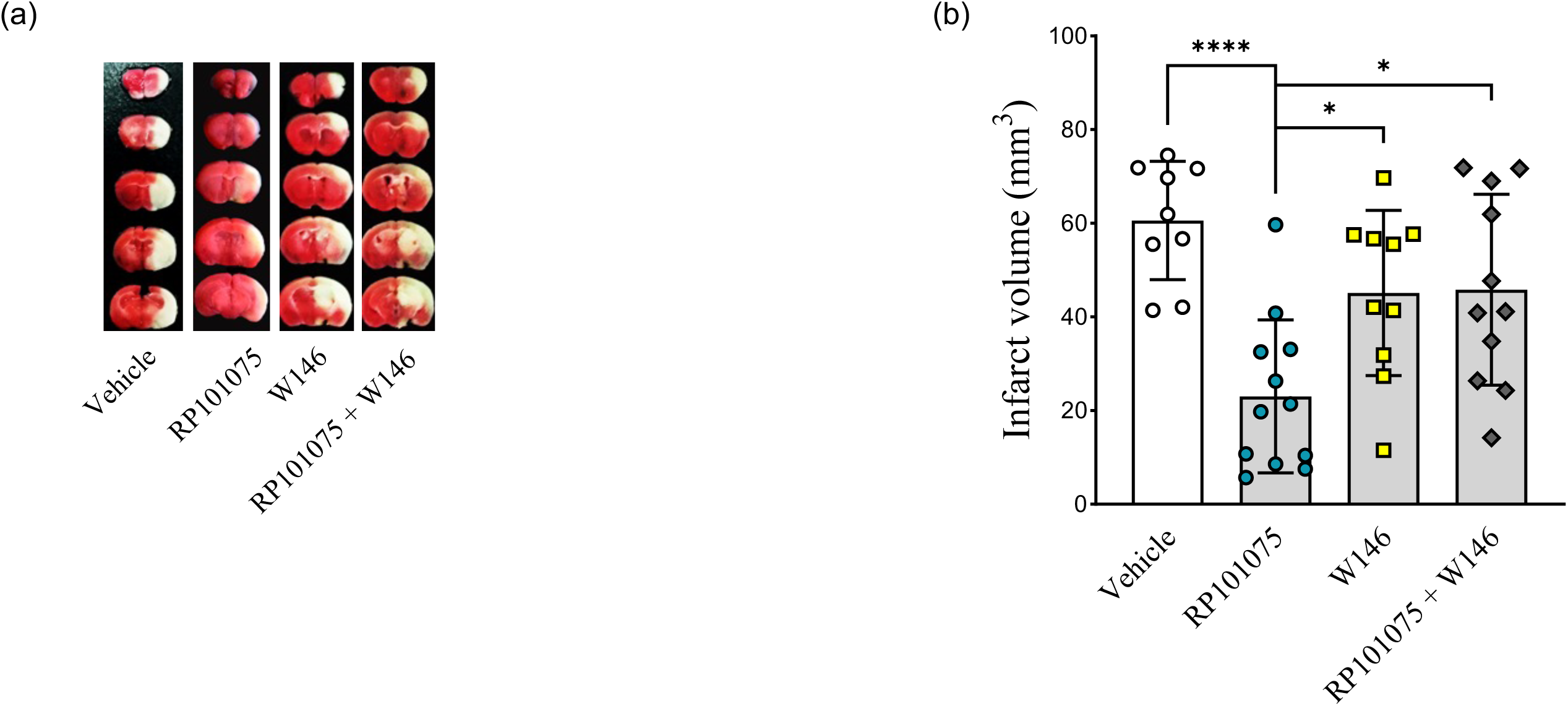
**Selective pharmacological blockade of S1PR1 reversed the reduction of brain infarct volume by RP101075 following tMCAO.** Infarct volume was assessed at day 1 via triphenyltetrazolium chloride (TTC) staining in mice treated with vehicle, RP101075 (0.6mg/kg), W146 (10μM) or RP101075 plus W146. Representative TTC stained slices are illustrated in panel (a). (b) Quantification of infarct volume assessed at day 1 after ischemia across treatment groups. Data are presented as mean ± SD; N = 9 in vehicle, 12 in RP101075, 10 in W146, 11 in RP101075 plus W146 (panel b). One-Way ANOVA test with Tukey post-hoc test was conducted to do compare the infarct volume. Vehicle vs RP101075 ****p < 0.0001, RP101075 vs W146 \**P* = 0.0228, RP101075 vs RP101075 + W146 \**P* = 0.0149.

### RP1010175 improves functional outcome post-tMCAO and this response is S1PR1 dependent

To evaluate functional outcome at recovery intervals on day 1, 3 and 7 after tMCAO, we used an 18-point mNSS score system that included behavioral tests (Figure 3). On day 1, there were no differences in scores between all groups. Mice receiving RP1010175 had significantly decreased mNSS scores compared to vehicle group on days 3 and 7 post occlusion, Day 3, 6.55 ± 1.015 vs 10.00 ± 0.57 (P=0.009); Day 7, 5.11 ± 0.77 vs 8.44 ± 0.89 (P=0.012). Selective pharmacological blockade using W146 attenuated RP101075’s protective response and returned mNSS scores back to those observed in tMCAO mice administered vehicle only RP101075+W146 vs vehicle: Day 1, 11.50 ± 0.94 vs 11.63 ± 0.69 (P=0.999); Day 3, 9.50 ± 0.62 vs 10.000 ± 0.58 (P=0.975); Day-7 8.17 ± 1.35 vs 8.44 ± 0.89 (P=0.995). W146 alone had no effect on neurologic improvement in tMCAO mice. These data suggest that post occlusion administration of RP101075 improves neurological outcome, and this response is S1PR1 dependent.

**Figure 3.**
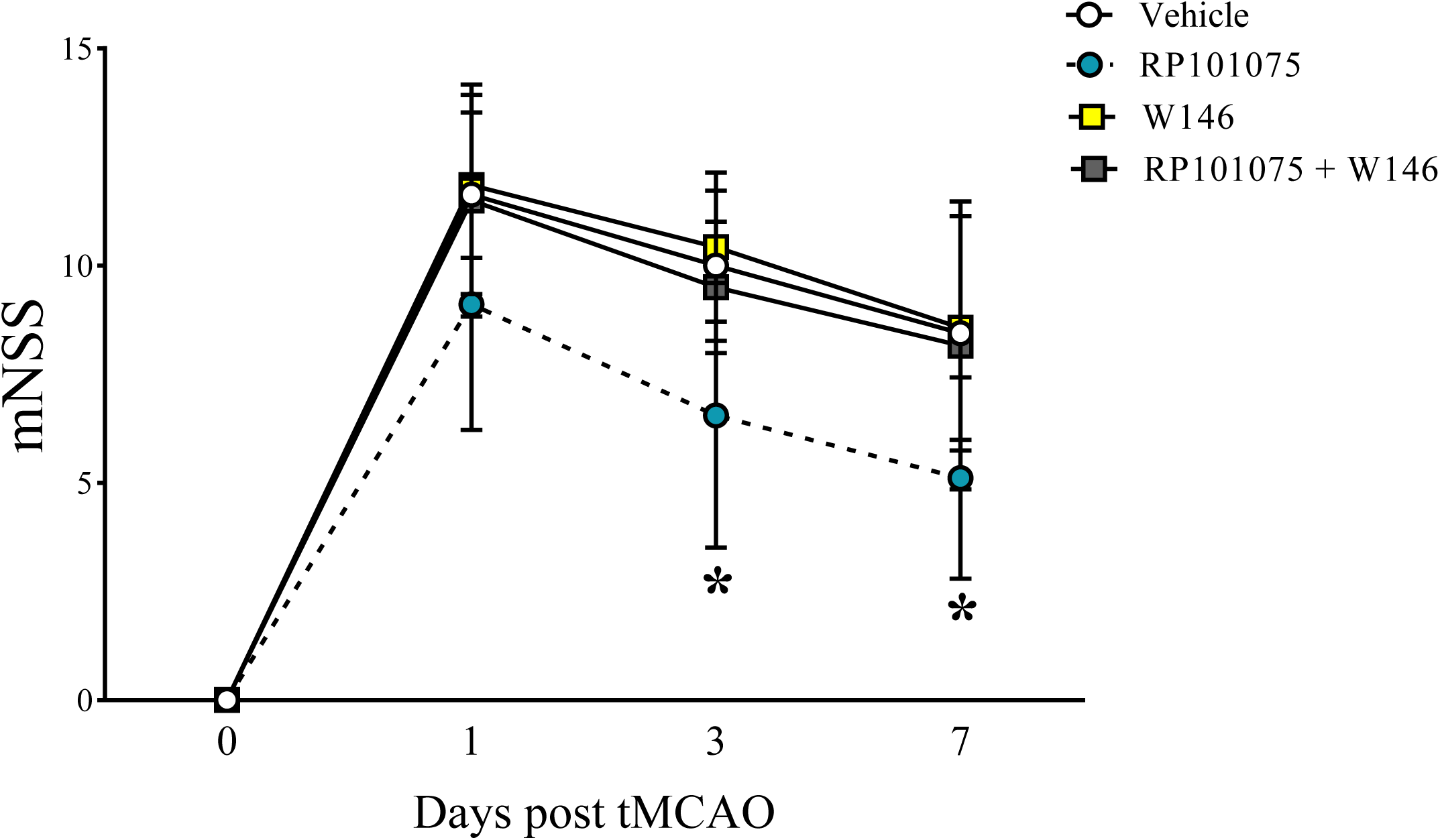
**RP101075 improved neurological function at days 3 and 7 following tMCAO and this response was S1PR1 dependent.** An 18-point modified neurological severity scores (mNSS) was implemented to evaluate neurological function in mice following either 1h tMCAO and treated with either vehicle, RP101075 (0.6mg/kg), W146 (10μM), or RP101075 plus W146 at days 0 (before MCAO) to 7 following reperfusion. Graphical illustration of mNSS scores is depicted and demonstrates a timeline of neurological severity in which worsened neurological function is given a greater mNSS score. A score of 13–18 indicated severe injury, 7–12 indicated moderate injury, and 1–6 indicated mild injury. Data are presented as mean ± SD; N = 9-11 in vehicle, 9 in RP101075, 7 in W146, and 6-8 in RP101075 plus W146. Two-Way ANOVA test with Tukey post-hoc test. Day 3 Vehicle vs. RP101075 \**P* = 0.0191, Day 7 Vehicle vs. RP101075 \**P* = 0.0249.

### Mouse whole brain S1PR1 protein levels increase following tMCAO and this response is region and endothelial cell specific

Since RP101075-mediated improvement in functional outcome demonstrated S1PR1 dependence, we assessed brain S1PR1 protein levels following tMCAO (Figure 4). 1 S1PR1 levels were measured in the ipsilateral and contralateral hemispheres and relative S1PR1 band intensities were normalized to the corresponding β-actin. Ipsilateral and contralateral hemispheres from the sham group revealed faint bands at ∼40 kDa indicating basal levels of S1PR1 receptor protein expression that was similar across hemispheric sections. Following tMCAO, a triplet band profile at 40-43 kDa for anti-S1PR1 increased in tissue collected in the ipsilateral hemisphere (Figure 4a). An approximate 2-fold increase in relative signal intensity of S1PR1 protein levels within the tMCAO ipsilateral hemisphere compared to sham operated controls was observed, tMCAO ipsilateral vs sham ipsilateral 3.792 ± 0.480 vs 1.557 ± 0.535 (P=0.0026). These data indicate that S1PR1 is upregulated following ischemia and 24h reperfusion. Moreover, this upregulation is regionally specific to the hemisphere relative to the focal infarct. Next, we characterized the S1PR1 protein expression profile of brain endothelium and resident CNS single cell suspensions following tMCAO using FACS. FACS analysis detected the relative intensity of S1PR1 in the tagged cell populations compared to the fluorescence minus one (FMO) internal control from each experiment (Figure 4b). S1PR1 was detected in all cell types interrogated, and in direct support of the previous western blot findings indicating a basal protein expression of S1PR1. Notably, comparisons between sham and tMCAO demonstrate a marked increase in S1PR1 protein expression within the brain endothelium, tMCAO vs Sham; 86.800 ± 1.640 vs 63.800 ± 1.308 (P<0.0001) (Figure 4c). These data suggest that the major alteration of S1PR1 levels following tMCAO in male mice is within the cerebrovascular endothelium.

**Figure 4.**
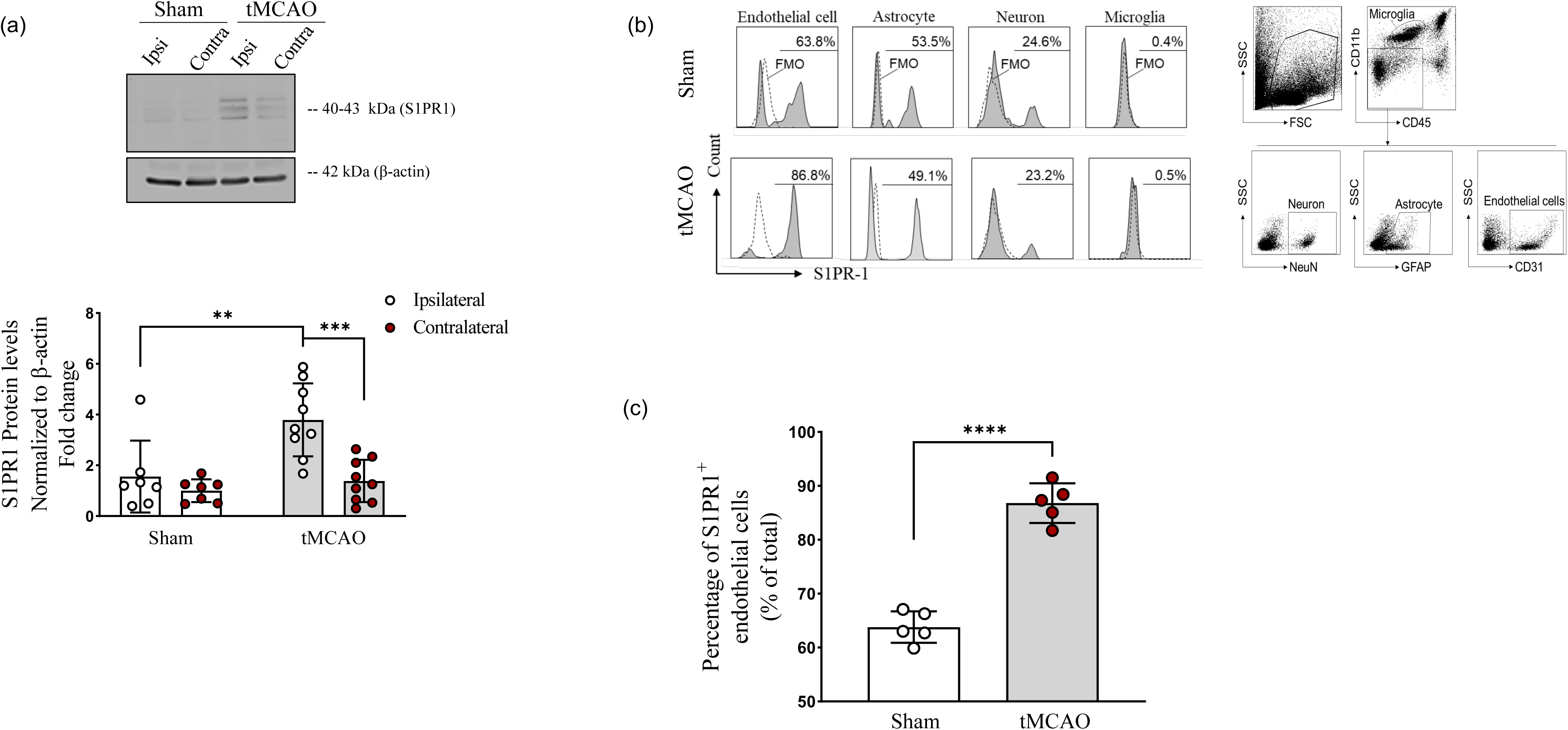
**S1PR1 protein levels increased in mouse brain endothelial cells following tMCAO.** Panel (a) illustrates a representative western blot of anti-S1PR1 and relative band intensities normalized to β-actin and averaged to the sham contralateral group values to calculate the S1PR1 expression fold changes in brain regions from ipsilateral sham and ipsilateral and contralateral tMCAO brain lysate. At 24h after 60 min tMCAO, cell suspensions were prepared from brain of male mice to run flow cytometry analysis. Gate strategy of flow cytometry analysis (panel b). Flow cytometry analysis determined the expression level (% of individual cell population) of S1PR1 cells in T (CD4^+^ or CD8^+^), NK (CD3^−^NK1.1^+^), neutrophil (CD11b^+^CD45hiLy6C^−^Ly6G^+^), astrocytes (GFAP^+^), endothelial cells (CD31^+^), microglia (CD11b^+^CD45^int^) and neuron (NeuN^+^) from sham and tMCAO brain lysate. A representative graph is shown in panel (c) which illustrates the percent of endothelial cells labeled with S1PR1 and demonstrated that tMCAO increased endothelial S1PR1 protein expression. Panel (a): N=7 in sham and N=9 in tMCAO for both contralateral and ipsilateral hemispheres; two-way ANOVA with Tukey’s multiple comparisons post hoc test; tMCAO ipsilateral vs sham ipsilateral \*\**P* = 0.0026, tMCAO ipsilateral vs. tMCAO contralateral \*\*\**P* = 0.0005; Panel (b): N=5/group; direct comparison was performed with unpaired *t* test with Welch’s correction; Sham vs. tMCAO \*\*\*\**P* < 0.0001. Data are presented as mean ± SD.

### Human brain endothelial cell health decreased and concomitant S1PR1 protein levels increased following HGD

Since murine endothelial S1PR1 levels were responsive to ischemic injury, we next addressed the impact HGD on brain endothelial S1PR1 protein levels (Figure 5). To validate our in vitro ischemic model, quantification of vesiculo-vacuolar like structures was assessed as an indicator of HGD-induced endothelial cell stress. We hypothesized these structures correlated to an endothelial structure referred to as the vesiculo-vacuolar organelle. This organelle, comprised of uncoated vesicles and vacuoles, has been previously shown to be within the cytoplasm of endothelial cells of tumor microvessels as well as normal venules and contribute to increased permeability.^46^ Visualization of HBMEC cytoplasmic vesiculo-vacuolar organelles (black arrows) via crystal violet staining following normoxia or HGD exposure is represented in Figure 5a and the quantification of the formations is illustrated graphically in Figure 5b. Basal amounts of vesiculo-vacuolar organelles were observed under normoxic conditions; however, HGD exposure induced a significant increase in vesiculo-vacuolar organelles per cell, Normoxia vs HGD; 3.048 ± 2.287 vs 13.48 ± 7.184 (P<0.0001) (Figure 5b). Additionally, our lab has previously observed that the same model of HGD decreased human brain vascular smooth muscle cell viability in a time-dependent manner from 3-9 h, ^27^ complimenting studies showing that oxygen-glucose deprivation as early as 2 h led to brain endothelial cell death.^47–49^ In this study, percent cell viability was determined via the ratio of live cell counts to total cell counts using trypan blue inclusion and are graphically illustrated in Figure 5c. In comparison to normoxic exposure at the 3h timepoint, HGD decreased cell viability, Normoxia vs HGD; 115.1 ± 6.494 vs 86.11 ± 6.538 (P<0.0001). Together, these data confirmed that our model of in vitro ischemia-like injury, HGD, was effective in inducing a decrease in HBMEC health. Next, following the validation of HGD we determined whether ischemia-like injury alters our target G-protein coupled receptor of interest in HBMECs. Following HGD (3h), S1PR1 protein levels increased (Figure 5d) Normoxia vs HGD; 0.95 ± 0.14 vs 1.43 ± 0.14 (P=0.0369). These data suggest, that like the protein expression profile we observed in the brain endothelium in our murine experimental stroke model, human brain endothelial cell S1PR1 also increases when cells are challenged with ischemia-like injury.

**Figure 5.**
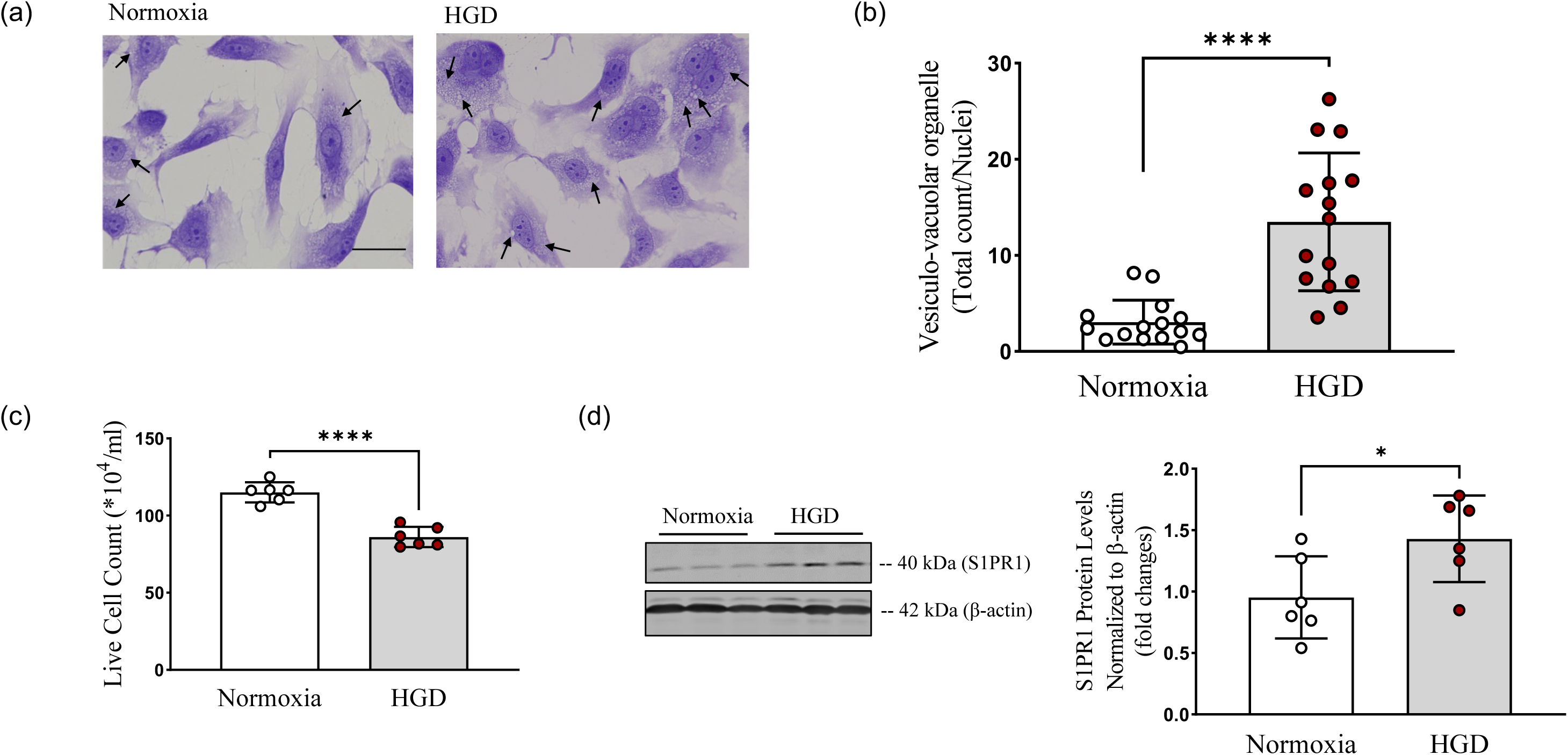
**HGD exposure decreased cell viability and increased S1PR1 levels in cultured human brain endothelial cells.** Prior to evaluating S1PR1 protein expression, we validated our in vitro HGD ischemia-like injury model by assessing cell health (quantification of vesiculo-vacuolar organelle formation) and viability following in vitro exposure to HGD. Panel (a) illustrates representative crystal violet images of human brain microvascular endothelial cells (HBMEC) exposed to either normoxia or HGD. Examples of HBMEC vesiculo-vacuolar organelles are highlighted by black arrows imaged at 40X magnification. Scale bar, 25μm. Bar graphs demonstrate the ratio of vesiculo-vacuolar organelle (panel b) quantitated normalized to the number of nuclei following normoxia or HGD for 3h. Panel (c) illustrates HBMEC viability expressed as the total number of live cells following either normoxia or HGD exposure. Panel (d) illustrates a representative western blot image of anti-S1PR1 and relative S1PR1 band intensities normalized to β-actin and averaged to the normoxia group to calculate the S1PR1 expression fold changes HGD endothelial cell lysate. Vesiculo-vacuolar organelle study: N=15 for vesicle independent samples; unpaired *t* test with Welch’s correction; normoxia vs. HGD \*\*\*\**P* < 0.0001. Live cell count studies are equal to N=6 independent samples; unpaired *t* test with Welch’s correction; normoxia vs. HGD \*\*\*\**P* < 0.0001. S1PR1 protein study: N=6 independent samples \**P* < 0.05. Data are mean ± SD.

### RP101075 and its precursor compound, ozanimod, both attenuated HGD-induced decreases in HBMEC health and this response was S1PR1 dependent

The proliferation of endothelial cells following ischemic injury is generally considered to begin after several days ^50, 51^ and is contributed to the secretion of multiple growth factors and subsequent receptor signaling; however, the acute proliferative response of HBMEC to ischemic injury has yet to be clarified. To determine the effect of selective S1PR1 targeting on HBMEC proliferation following HGD exposure, BrdU staining and live cell counts were measured. An RP101075 dose response for cell proliferation in our cell model exposed HGD for 3h was first determined (Figure 6a). The proliferative status of HBMECs exposed to HGD plus vehicle served as the control and remained unaltered with treatment of RP101075 at doses of 1nM and 5nM but was significantly increased following 10nM treatment, HGD + Vehicle vs HGD + RP101075 10nM; 16.06 ± 3.236 vs 30.73 ± 4.472 (P=0.0412). During normoxic conditions, RP101075 had no effect; however, 10nM RP101075 reversed the HGD-induced decrease in cell proliferation, Normoxia + Vehicle vs HGD + Vehicle; 29.13 ± 1.073 vs 16.06 ± 3.236 (P=0.0209) (Figure 6b). These data suggest that HGD exposure at an acute time point of 3h induces a decrease in endothelial cell proliferation and treatment with a selective S1PR1 ligand reverses this response. Moreover, HBMECs, when treated with RP101075 under HGD exposure, returned to approximately to the observed normoxic baseline and did not exceed this level, which suggests a potential return to replacement capability rather than aberrant and uncontrolled proliferation to improve vascular integrity in response to an acute ischemic injury.

**Figure 6.**
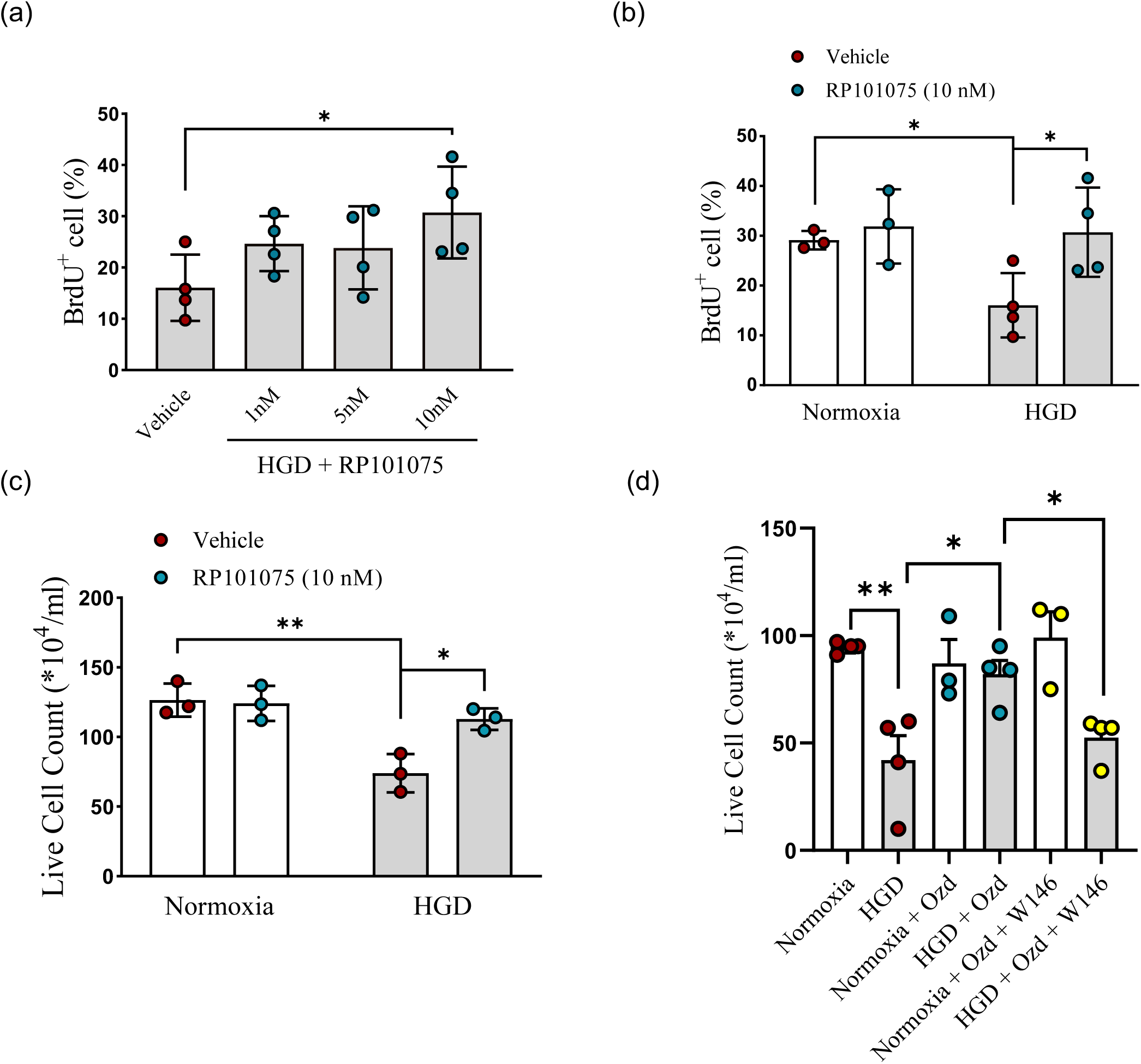
**Selective S1PR1 ligands increased cultured human brain endothelial cell proliferation and live cell count during HGD.** HBMEC proliferation was determined via bromodeoxyuridine (BrdU) staining and subsequent flow cytometric analysis. Three doses of RP101075 were administered (1nM, 5nM, 10nM) (panel a). The percentage of endothelial cells positive for BrdU staining are represented in a bar graph as a percentage of BrdU positive cells relative to the remaining unstained cells within the same individual cohort loaded and analyzed. Panel (b) illustrates a bar graph which depicts the relative percentage of BrdU positive HBMECs following exposure to either normoxia or HGD for 3h and treatment with either RP101075 (10nM) or vehicle. Trypan blue staining of HBMECs exposed to either normoxia or HGD for 3h and treated with either RP101075 (10nM) (panel c) or ozanimod (Ozd; 0.5nM), W146 (selective S1PR1 antagonist; 10 μ M), or associated vehicles (panel d) was used to determine live cell count. Panel (a): N=4 independent samples; direct comparison was performed with unpaired *t* test with Welch’s correction; HGD + Vehicle vs. HGD + RP101075 (10nM) \**P* = 0.0412. Panel (b): N=3-4 independent samples; direct comparisons were performed with unpaired *t* test with Welch’s correction; Normoxia + Vehicle vs. HGD + Vehicle \**P* = 0.0209; HGD + Vehicle vs. HGD + RP101075 (10nM) \**P* = 0.0412. Panel (c): N=3-4 independent samples; direct comparisons were performed with unpaired *t* test with Welch’s correction; Normoxia + Vehicle vs. HGD + Vehicle \*\**P* = 0.0075; HGD + Vehicle vs. HGD + RP101075 (10nM) \**P* = 0.0131. Panel (d): N=3-4 independent samples; direct comparisons were performed with unpaired *t* test with Welch’s correction; Normoxia + Vehicle vs. HGD + Vehicle \*\**P* = 0.0039. HGD + Vehicle vs. HGD + Ozd (0.5nM) \**P* = 0.0229. HGD + Ozd vs HGD + Ozd + W146 \**P* = 0.0121. Data are mean ± SD.

Although it has been reported that endothelial cell death contributes to vascular remodeling under physiological brain development [Reviewed in ^52^], the continued integrity of cerebrovascular function following ischemic stroke is in part dependent on the viability of the endothelial cells. [Reviewed in ^53^] To this end we sought to expand upon our previous proliferation observations by next examining the viability of HBMECs as a potential target of vascular integrity. In line with the previous BrdU staining data, there was a decrease in HBMEC viability following 3h HGD exposure compared to normoxic controls measured via trypan blue staining, Normoxia + Vehicle vs HGD + Vehicle; 126.5 ± 6.874 vs 74.00 ± 7.943 (P=0.0075) (figure 6c). When treated with RP101075, normoxic controls exhibited no change but there was a significant attenuation of the HGD-mediated decrease in HBMEC viability, HGD + Vehicle vs HGD + RP101075; 74.00 ± 7.943 vs 112.8 ± 4.512 (P=0.0131) (figure 6c). In addition to the RP compound, we tested the effect of its parent compound, ozanimod, to test whether an additional selective S1PR1 ligand could attenuate HGD-induced decreases in endothelial cell viability, and we further tested whether this response was S1PR1 dependent. Consistent with the previous experiment, we observed a significant decrease in HBMEC viability following 3h HGD exposure compared to normoxic controls and treatment with ozanimod significantly attenuated this response, HGD + Vehicle vs HGD + Ozd; 42.00 ± 11.45 vs 82.00 ± 6.494 (P=0.0229) (figure 6d). The inhibitor, W146, had no effect in the normoxic control group; however, W146 reversed ozanimod’s response during HGD, HGD + Ozd vs HGD + Ozd + W146; 82.00 ± 6.494 vs 52.50 ± 5.188 (P=0.0121) (Figure 6d). These data suggest that S1PR1 ligands via S1PR1 enhance endothelial cell survival during acute ischemia-like injury.

### Selective S1PR1 ligand attenuated both MMP-9 and MMP-2 activity and preserved claudin-5 levels following HGD/R injury

MMP-9 as well as MMP-2 have been implicated in the pathogenic ictus of acute ischemic stroke as it pertains to increased permeability at the blood brain barrier, ^54–57^ which can lead to worsened outcomes.^58^ In efforts to elucidate a potential additional mechanism by which selective S1PR1 ligands reduced infarct volume and improved neurological function we investigated the impact of ozanimod, the parent compound of RP101075, on HBMEC derived MMP-9 and MMP-2 activity (figure 7a). Utilizing a model of in vitro ischemia-reperfusion like injury, HGD/R, ozanimod decreased both MMP-9, HGD/R + Vehicle vs HGD/R + Ozd (0.5nM); 1.000 ± 0.01075 vs 0.7076 ± 0.2019 (P=0.0276) (figure 7b) and MMP-2, HGD/R + Vehicle vs HGD/R + Ozd (0.5nM); 1.000 ± 0.2957 vs 0.4675 ± 0.08461 (P=0.0134) (figure 7c) activity at 24h of in vitro reperfusion when treated with ozanimod immediately following 3h HGD exposure. These responses were then both reversed when treated with selective S1PR1 antagonist, W146, immediately at the point of reperfusion but 0.25h prior to ozanimod treatment, HGD/R + Ozd (0.5nM) vs HGD/R + Ozd (0.5nM) + W146 (10μM); 0.7076 ± 0.2019 vs 2.794 ± 1.542 (P=0.0364) (figure 7b), HGD/R + Ozd (0.5nM) vs HGD/R + Ozd (0.5nM) + W146 (10μ M); 0.4675 ± 0.08461 vs 1.543 ± 0.7735 (P=0.0327) (figure 7c). Additionally, we examined levels of claudin-5 mRNA in the HBMECs following HGD/R both at 12h and 24h post-ischemic like injury, as claudin-5 has been referred to previously as the gatekeeper of neurological function.^59^ There was no observable change in claudin-5 mRNA levels at 12h post-injury; however, unexpectedly we observed a decrease in claudin-5 in HBMECs treated with ozanimod following HGD/R (3h/12h), HGD/R + Vehicle vs HGD/R + Ozd (0.5nM); 0.9015 ± 0.1798 vs 0.4622 ± 0.1069 (P=0.0057) (figure 7d). When examining levels at 24h post-injury and in concordance with our in vivo observations, there was a decrease in claudin-5 mRNA, Normoxia + Vehicle vs HGD/R + Vehicle; 1.049 ± 0.3257 vs 0.7576 ± 0.2028 (P=0.0497) (figure 7e). This decrease of claudin-5 following HGD/R (3h/24h) was attenuated by ozanimod, HGD/R + Vehicle vs HGD/R + Ozd (0.5nM); 0.7576 ± 0.2028 vs 1.074 ± 0.3154 (P=0.0315) (figure 7e). Similarly, following HGD/R (3h/24h) there was a notable decrease in zonula occludens 1 (ZO-1) mRNA, Normoxia + Vehicle vs HGD/R + Vehicle; 1.012 ± 0.1595 vs 0.6788 ± 0.1967 (P=0.0023) (figure 7f). However, unlike claudin-5, treatment with ozanimod had no impact on ZO-1 mRNA levels following exposure to HGD/R (3h/24h). These data suggest that S1PR1 ligands in part via S1PR1 provide barrier protection at the level of the endothelium at 24h of reperfusion following an acute ischemia-like injury.

**Figure 7.**
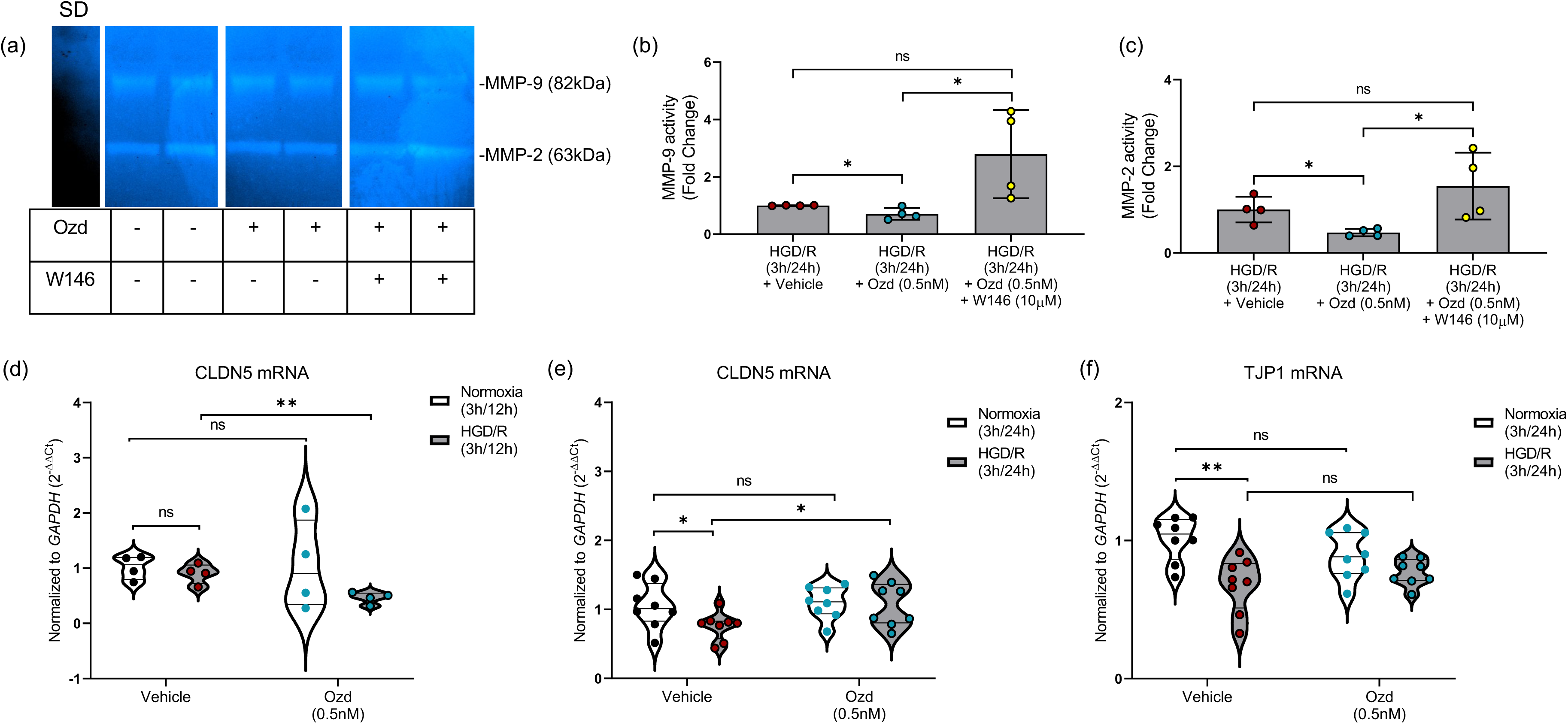
**S1PR1 selective ligand decreased extracellular MMP-9 and MMP-2 activity and preserved claudin-5 mRNA levels.** Panel (a) illustrates standard gelatin zymography which was utilized to visualize the band migration and quantitate enzymatic activity for two distinct matrix metalloproteinase bands, MMP-9 (82kDa) and MMP-2 (63kDa), in conditioned media collected from HBMECs following treatment with either vehicle, ozanimod (Ozd; 0.5nM), or W146 (selective S1PR1 antagonist; 10μM), and exposure to HGD/R for 3h/24h. Standard, SD. Bar graphs depict MMP-9 (panel b) and MMP-2 (panel c) activity by inverse densiometric analysis and expressed as fold change ± SD. Panel (d) depicts qRT-PCR mediated quantitation of *CLDN5* mRNA present within HBMECs exposed to either normoxia or HGD/R (3h/12h) and treated with either vehicle or ozanimod (Ozd; 0.5nM) normalized to the housekeeping gene (*GAPDH*) and expressed as 2^−ΔΔ*Ct*^. Panels (e) and (f) depict qRT-PCR mediated quantitation of *CLDN5* (panel e) and *TJP1* (panel f) mRNA present within HBMECs exposed to either normoxia or HGD/R (3h/24h) and treated with either vehicle or ozanimod (Ozd; 0.5nM) normalized to the housekeeping gene (*GAPDH*) and expressed as 2^−ΔΔ*Ct*^. Panel (b): N=4 independent samples; planned direct comparisons were performed with unpaired *t* test. HGD/R (3h/24h) + Vehicle vs. HGD/R (3h/24h) + Ozd (0.5nM) \**P* = 0.0276, HGD/R (3h/24h) + Ozd (0.5nM) vs. HGD/R (3h/24h) + Ozd (0.5nM) + W146 (10μM) \**P* = 0.0364, HGD/R (3h/24h) + Vehicle vs. HGD/R (3h/24h) + Ozd (0.5nM) + W146 (10μM) *P* = 0.0589. Panel (c): N=4 independent samples; planned direct comparisons were performed with unpaired *t* test. HGD/R (3h/24h) + Vehicle vs. HGD/R (3h/24h) + Ozd (0.5nM) \**P* = 0.0134, HGD/R (3h/24h) + Ozd (0.5nM) vs. HGD/R (3h/24h) + Ozd (0.5nM) + W146 (10μM) \**P* = 0.0327, HGD/R (3h/24h) + Vehicle vs. HGD/R (3h/24h) + Ozd (0.5nM) + W146 (10μM) *P* = 0.2378. Panel (d): N=4 independent samples; HGD/R (3h/12h) + Vehicle vs. HGD/R (3h/12h) + Ozd (0.5nM) *P* =0.0057. Panel (e): N=8 independent samples; Normoxia + Vehicle vs. HGD/R (3h/24h) + Vehicle \**P* = 0.0497, HGD/R (3h/24h) + Vehicle vs. HGD/R (3h/12h) + Ozd (0.5nM) \**P* = 0.0315. Panel (f): N=8 independent samples; Normoxia + Vehicle vs. HGD/R (3h/24h) + Vehicle \*\**P* = 0.0023. Data are mean ± SD.

## Discussion

In this study, we tested a novel selective S1PR1 ligand, RP101075, and to an extent its parent compound ozanimod, as pharmacologic tools to help identify a clinically relevant approach to stroke treatment with the potential to involve the brain endothelium. Our findings suggest that RP101075 administered post occlusion and during reperfusion affords neuroprotection and improves neurologic outcome via S1PR1 activation following tMCAO. Additionally, we observed that RP101075 enhances endothelial cell survival and proliferation during HGD. Moreover, targeting selectively targeting S1PR1 within HBMECs decreased MMP-9 and MMP-2 as well as preserved claudin-5 levels. Concomitantly, we observed that basal brain S1PR1 protein levels increase following tMCAO, specifically in the ischemic ipsilateral hemisphere, and levels are greatest in CNS endothelial cells compared to other CNS cell types. Similar to our in vivo tMCAO model, basal S1PR1 protein levels in HBMECs increase following HGD. Together, these findings suggest that RP101075 via S1PR1 activation provides protection against cerebral ischemic insult and this protection potentially involves preservation of endothelial cell health and enhanced barrier properties.

In our in vivo studies, the most compelling findings are that RP101075 administration at onset of reperfusion reduced infarct volumes and improved neurologic scores following tMCAO. Reduction in brain injury in our male mouse stroke model is in agreement with a previous study using a milder form of tMCAO (45 min occlusion). Brait et al ^20^ reported that LASW1238, another selective S1PR1 ligand, as well as fingolimod (non-selective S1PR modulator) administered post occlusion elicited a reduction in infarct volume. However, unlike our findings demonstrating that RP101075 improved neurologic outcome, they reported no effect of their test compounds on neurologic function. The possible discrepancy may involve differences in dose administration and/or length of occlusion used in each model. For example, Brait et al (2016) subjected their mice to 45 minutes of occlusion/24h reperfusion whereas in our studies we challenged mice with 60 min occlusion/24h reperfusion. Although they observed reduced infarct volumes using 10 mg/kg for both LASW1238 and fingolimod, in our studies we observed an efficacious response using a lower concentration of RP101075 (0.6 mg/kg). We also observed that selective inhibition of S1PR1 reversed both the beneficial effect of RP101075 on injury and neuroprotection. Future studies are planned to determine if RP101075s parent compound, ozanimod, elicits a similar neuroprotective efficacy.

Clinically, groundbreaking treatments like thrombolysis and mechanical clot retrieval are effective^60–62^; however, these interventions are associated with high mortality risk and limit options for AIS patients as delayed hospital arrival or lack of access to a stroke center remain a concern. Fingolimod demonstrated promising therapeutic potential for AIS treatment, however due to its broad S1PR pharmacological actions, cardiac safety remains a concern.^63, 64^ Studies have reported that concerns for its cardiac safety involved activation of S1PR3. In adult naive male mice, fingolimod produced a dose-dependent transient reduction in heart rate, and this response was S1PR3 dependent. ^65, 66^ However, unlike fingolimod, RP101075 has been reported to have almost no affinity to S1PR3.^29^ It has been reported that RP101075 has affinity to both S1PR1 and S1PR5 with the half maximal effective concentration (EC50) of RP101075 reported to be more than 20 times less for S1PR1 (0.27nM±0.06) than for S1PR5 (5.9nM±1.0) ^67^, which indicates that RP101075 has a significantly higher affinity to S1PR1 than S1PR5. Moreover, S1PR5 is predominantly reported in the white matter tracts and oligodendrocytes.^68^ We confirmed the involvement of S1PR1 rather than S1PR5 by treating with W146. W146 is a highly selective antagonist for S1PR1 ^69^ and at a concentration of 10 μM has been reported to elicit no off-target S1PR (i.e. S1PR2, S1PR3, and S1PR5) agonist nor antagonist effects.^40^ W146 abolished the beneficial effects of RP101075 on brain injury and outcome following tMCAO, suggesting that S1PR1 rather than S1PR5 plays the vital roles in RP101075 treatment.

Our finding of increased S1PR1 protein level expression in this study following acute ischemic injury was consistent with previous studies reporting upregulation of *Edg-1* gene 1h after brain ischemia injury.^70^ However, relatively few studies have characterized S1PR1 protein levels/expression profiles under pathophysiological conditions related to stroke. For instance, a study using a common carotid artery occlusion method of ischemic injury observed a temporal increase in overall S1PR1 mRNA levels in the temporo-parietal cortex which included leptomeningeal arteries at days 4 and 7 after CCAO.^71^ In the same study authors also observed a colocalization of S1PR1 expression to CD31 and showed an increase in S1PR1 labeled endothelial cells quantified as the percentage of CD31 labeled endothelial cells as an area (representing a peak ∼45% of endothelial cells) labeled with S1PR1 at days 1-14. In a separate study utilizing a 2h tMCAO method of ischemic injury, there was observed immunofluorescence staining of S1P1 expressed in neurons within the periinfarct cortex at 24 hours after MCAO; however, no quantification of S1PR1 expression across treatment groups were performed.^72^ Using in vivo kainic acid administration or traumatic brain injury model, studies have reported an increase in S1PR1 expression in neural stem cells and hippocampal neurons respectively.^73, 74^ Our results provide further insight to the spatial regulation of S1PR1 alteration post-ischemia at the protein level and was further interrogated at the level of the individual cell within the brain following ischemia injury via flow cytometry. Matsumoto et al. 2020 reported an increase in S1PR1 in the peri-infarcted lesion following tMCAO 24h.^75^ However, to the best of our knowledge this is the first report of differential S1PR1 protein expression changes in CNS types following focal ischemic injury which demonstrates the most pronounced alteration in S1PR1 is within the brain endothelium.

HBMECs were included in the study design to confirm ischemia-induced expression alternations in a human cell derived model. In a broader context, targeting S1PR1 within the endothelium following an acute ischemic injury is an attractive therapeutic target. Although the size and position of the lesion may vary in patient presentation, the salvageable penumbra may envelope a number of penetrating small arteries and arterioles, and their associated capillaries, which are predominated by endothelial cells. Thus, within the penumbra, the extent of vascular injury may be greatest, leading to acute endothelial dysfunction. Endothelial dysfunction disrupts the mechanism of vascular homeostasis regulation, predisposing the vessel wall to vasoconstriction, leukocyte adhesion, platelet activation, oxidative stress, thrombosis, coagulation, and inflammation. While the underlying physiological relevance of the S1PR1 upregulation is unclear, further investigation into endothelial sphingolipid metabolism, although beyond the scope of this study, may provide a better understanding. In this study we postulated that this region-specific concentration may be a physiological response to the endothelial dysfunction in the penumbra surrounding the necrotic core. Considering our finding describing the neuroprotective effects of RP101075 at an acute time point, the ligand independent upregulation of S1PR1 may be a protective response to damage. However, deeper examination into the cellular regulation of S1PR1 expression in the context of ischemia as well as presence of selective ligands is warranted.

Cerebral endothelial cells are the foundation of the BBB. Loss of endothelial cell after ischemia permits penetration of intravascular proteins, fluid, and peripheral immune cells into the extracellular space, resulting in vasoactive edema and expansion of tissue damage.^76^ However, some papers reported that early stage endothelial cell damage is usually not accompanied by detachment of endothelial cells from the vessel wall; reflecting subtler changes, the adjacent endothelial cells fuse together by intercellular junctions anchored to the actin cytoskeleton.^77^ Actin functions to sustain the shape of endothelial, under HGD stress, actin filaments are polymerized into linear stress fibers, which result in morphological changes and impair junctional sealing efficiency.^78–80^ Our study revealed that RP101075 reversed the HGD-induced loss of endothelial cell viability and proliferation, which suggests that the protective effect of RP101075 on endothelial cells may contribute to maintaining BBB integrity. This is further substantiated by our findings that the parent compound, ozanimod, attenuated HBMEC derived MMP-9 and MMP-2 activity at 24h following 3h ischemic-like injury in an S1PR1 dependent manner. Further propagating the therapeutic potential of selectively targeting endothelial S1PR1 to promote barrier integrity within the cerebrovasculature. In addition to attenuated MMP activity, we also observed that the same selective S1PR1 ligand at the same time point, increased HBMEC claudin-5 mRNA levels which substantiates the potential barrier protective properties of endothelial S1PR1. These data compliment preliminary findings from our lab which demonstrate that 3h HGD exposure concomitantly increased vascular cell adhesion molecule 1 and decreased claudin-5 expression in HBMECs and treatment with the selective S1PR1 ligand, ozanimod, attenuated both responses (Wendt and Gonzales, unpublished observation). These data suggest that S1PR1 activation may also play a role in reducing endothelial cell adhesion molecule expression in addition to rescuing tight junction protein levels. This notion is supported by various mechanistic studies describing that S1PR1 signaling in endothelial cells leads to activation of the Rac pathways, stabilizing the cytoskeleton and enhancing cell-cell contact through adherens and tight junctions on the cell membrane and stabilizes cytoskeletal integrity.^81^ Further studies may include the efficacy of RP101075 on HGD induced F-actin, adhesion molecule, and intercellular junction alterations to clarify the mechanism of RP101075 in preservation of BBB integrity.

In summary, we addressed the efficacy of RP101075 on ischemic stroke and specifically characterized the S1PR1 expression alteration following ischemic injury at the level of endothelial cells. We also observed the protective effect of RP101075, and ozanimod (parent compound) on endothelial cells during HGD. We acknowledge that the complex pathophysiological cascade shown to occur during AIS cannot be exactly modeled in an in vitro setting and we acknowledge that the observations made in vivo and in culture conditions are parallel, but are not casually linked. However, in vitro studies using human primary cells do allow the investigation of specific basic cellular and molecular mechanisms under conditions of hypoxia or oxygen and glucose deprivation which is similar to what is observed during AIS. Based on our results in this study we posit that selective S1PR1 ligands are in part working at the level of MMP-9, MMP-2, and claudin-5 to attenuate ischemia-induced decreases in HBMEC barrier integrity which correlate to our observations made in vivo of reduced infarct volume followed by improved neurological function. However, due to the complex and interconnectivity of GPCRs, it is also likely that S1PR1 elicits multiple responses to attenuate ischemic stroke-like associated pathologies related to endothelial barrier function. Thusly, future studies are planned to investigate specific mechanism(s) of action in the cerebrovasculature in vivo including modulation of Akt, PI3K, Rho, and upstream mediators as well as barrier proteins in freshly dissected, uncultured, endothelial cells. Our results suggest that S1PR1 ligand may serve as important beneficial modulators against ischemic damage and their protective effects on brain vascular endothelial cells lay a foundation for further studying the mechanism(s) of selective ligands like RP101075 and or ozanimod against ischemic brain injury, especially at the cerebrovascular endothelial level.

## Acknowledgements

Funding support by a Valley Research Partnership P2 Award (RJG) [VRP37], the American Heart Association (QL) [16SDG27250236] and (RJG) [19AIREA34480018] Awards. Thank you Robert J. Peach and Paul A. Frohna (Receptos Inc. San Diego) for providing RP100175 as a gift.

## Author’s contributions

RJG and QL conceived and designed this study. SS, YL, TW and RJG wrote the paper. SS, YJL, TW, KS, WJ and RJG performed the experiments; SS, YJL, TW and RJG analyzed the data; SS, YJL, TW, QL and RJG interpreted the results of experiments; SS, YJL, TW QL and RJG prepared the figures and drafted/edited the manuscript.

